# ZRC3308 monoclonal antibody cocktail shows protective efficacy in Syrian hamsters against SARS-CoV-2 infection

**DOI:** 10.1101/2021.09.16.460724

**Authors:** Pragya D Yadav, Sanjeev Kumar Mendiratta, Sreelekshmy Mohandas, Arun K Singh, Priya Abraham, Anita Shete, Sanjay Bandyopadhyay, Sanjay Kumar, Aashini Parikh, Pankaj Kalita, Vibhuti Sharma, Hardik Pandya, Chirag G Patel, Mihir Patel, Swagat Soni, Suresh Giri, Mukul Jain

## Abstract

We have developed a monoclonal antibody (mAb) cocktail (ZRC-3308) comprising of ZRC3308-A7 and ZRC3308-B10 in the ratio 1:1 for COVID-19 treatment. The mAbs were designed to have reduced immune effector functions and increased circulation half-life. mAbs showed good binding affinities to non-competing epitopes on RBD of SARS-CoV-2 spike protein and were found neutralizing SARS-CoV-2 variants B.1, B.1.1.7, B.1.351, B.1.617.2 and B.1.617.2 AY.1 *in vitro*. The mAb cocktail demonstrated effective prophylactic and therapeutic activity against SARS-CoV-2 infection in Syrian hamsters. The antibody cocktail appears to be a promising candidate for the prophylactic use and for therapy in early COVID-19 cases which have not progressed to severe disease.

## Introduction

Since the first report of occurrence of SARS-COV-2 infection in China on December 30, 2019, the virus has spread rapidly worldwide, accounting for more than 200 million cases and 4 million deaths as on 9^th^ August 2021^1^. Even though, more than twenty vaccines have been granted Emergency Use Authorization (EUA) in multiple countries, vaccination rate is skewed globally due to limited accessibility to low- and middle-income countries^2,3^. Due to the inequitable access of vaccines and emergence of new variants having immune escape property, the development of herd immunity may take years^3^. Therefore, there remains an unfulfilled need for therapeutic agents to restrict the morbidity and mortality. Convalescent plasma transfusion has been used as therapeutic option in case of novel viral diseases like Ebola virus disease, SARS, MERS and coronavirus disease 2019 (COVID-19) but the variability in the donor antibody titers and risk of blood borne diseases remained as hurdles ^4–7^. Monoclonal antibody (mAb) based therapy became an alternative to overcome the limitations of convalescent plasma therapy. The favorable safety profiles and comparatively lesser time for generation and approval made mAbs a feasible therapeutic alternative in case of many emerging disease threats.

Monoclonal antibodies can serve as an adjunct to the prophylactic strategy for COVID-19 infections in high-risk groups like aged and immunocompromised people who have suboptimal responses to vaccination^8^. The techniques of combinatorial display libraries, humanized mice and B cell isolation methods have aided in rapid recovery of many antiviral mAbs^9^. Palivizumab was the first mAb to be approved for treatment against Respiratory Syncitial Virus in 1998^10^. In 2020, a combination of 3 mAbs, atoltivimab, maftivimab, and odesivimab-ebgn were approved by United States Food and Drug Administration (USFDA) for Zaire Ebola virus therapy ^11,12^. More than 200 research laboratories across world are working on developing highly potent recombinant human mAbs against SARS-CoV-2 to provide unlimited supply of high-quality product to patients. USFDA has granted EUA for few of the anti-SARS-CoV-2 mAbs targeting epitopes on receptor binding domain (RBD) of the S protein like bamlanivimab plus etesevimab, casirivimab plus imdevimab and sotrovimab for treatment of mild to moderate COVID-19 ^13–15^. We have developed a cocktail (ZRC-3308) of two highly potent, neutralizing, humanized mAbs, ZRC3308-A7 (CAS RN: 2640223-84-1) and ZRC3308-B10 (CAS RN: 2640224-48-0) that bind with nanomolar affinities to non-competing epitopes on the RBD of the spike protein of SARS-CoV-2. Both of these mAbs were designed to have reduced immune effector functions and increased circulation half-life. Here we describe the *in vitro* biological properties of the mAbs and *in vivo* evaluation against SARS-CoV-2 infection in Syrian hamster model.

## RESULTS

### Binding of ZRC3308 mAbs to RBD and Spike protein Trimer

ZRC-3308 cocktail showed similar binding profiles to RBD and S Trimer protein when compared with its individual components, ZRC3308-A7 and ZRC3308-B10 (Fig.1a-1f). The binding to both RBD and S trimer protein were with high affinities in the sub nM ranges for both the individual mAbs and their cocktail (Table 1). Pair-wise epitope binning experiment was performed to determine whether each of the two mAbs of the cocktail would bind to RBD even in the presence of the other one. The binding of each antibody to RBD was assessed after the other mAb had been allowed to bind first. Binding of ZRC3308-B10 was observed on ZRC3308-A7 precaptured channels and *vice versa* indicating that both the antibodies bind to distinct epitopes on RBD of spike protein as depicted in sensograms (Fig. 1g, 1h).

**Fig. 1:**
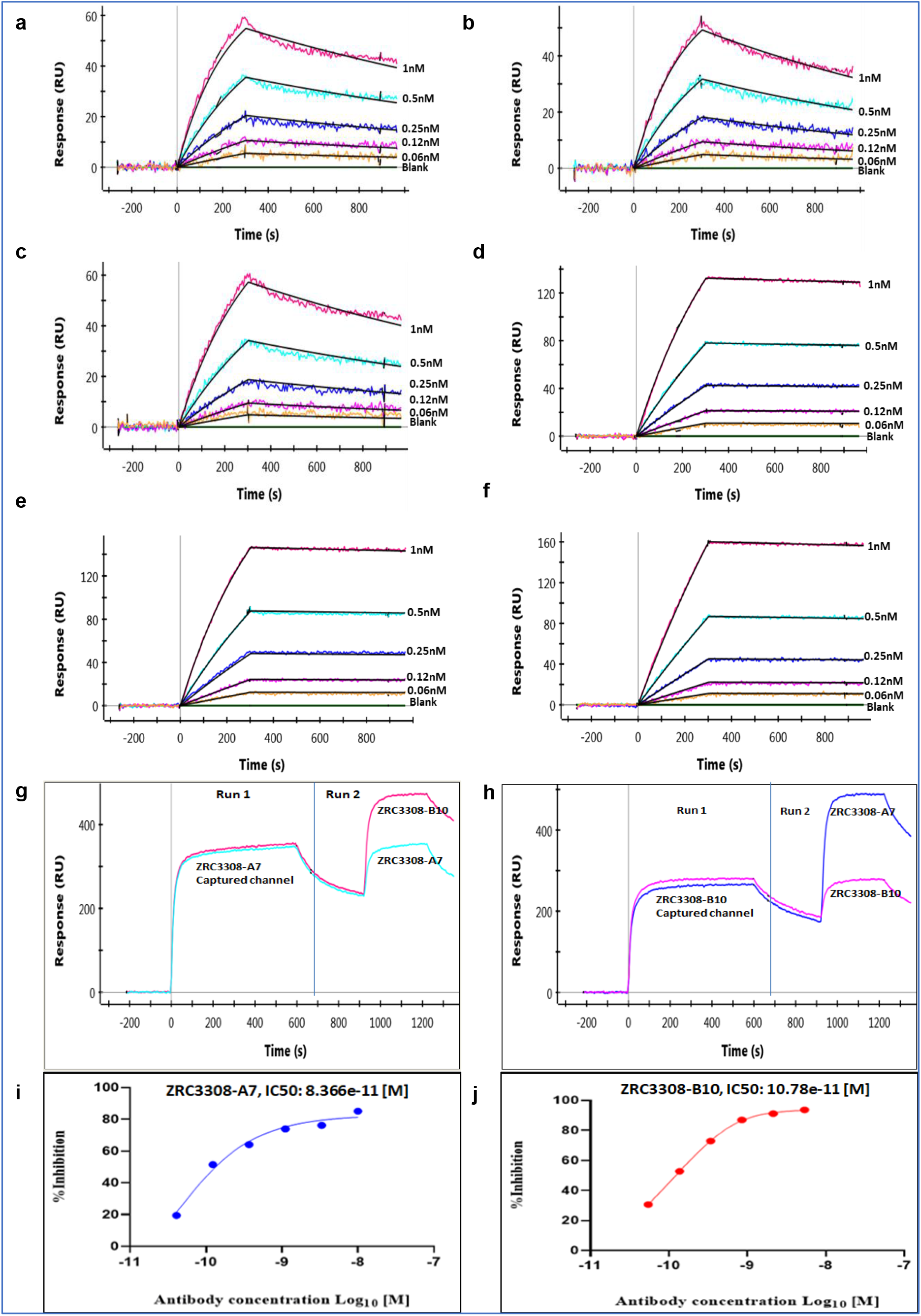
Binding of ZRC3308 to RBD and Spike protein trimer. Binding sensograms of **a.** ZRC3308-A7 **b.** ZRC3308B10 and **c.** ZRC3308 cocktail to RBD protein and **d.** ZRC3308-A7 **e.** ZRC3308-B10 and **f.** ZRC3308 cocktail to S protein. **g.** Binding of ZRC3308-B10 on ZRC3308-A7 precaptured channels and **h.** binding of ZRC3308-A7 on ZRC3308-B10 precaptured channels. Inhibition of RBD binding to ACE2 in presence of **i.** ZRC3308-A7 and **j.** ZRC3308-B10.

**Table 1.** Binding of ZRC3308 to RBD, S trimer protein, rhFcRn and FcγRIIIa-Phe. The kinetic rate constants of ZRC3308-A7, ZRC3308-B10 and ZRC3308 cocktail binding.

| Antibody | RBD |  |  | S trimer |  |  |
| --- | --- | --- | --- | --- | --- | --- |
|  | Ka | kd | KD | Ka | kd | KD |
|  | 1/Ms | 1/s | M | 1/Ms | 1/s | M |
| ZRC3308-A7 mAb | 4.57E+06 | 5.42E-04 | 1.19E-10 | 2.80E+06 | 4.53E-05 | 1.64E-11 |
| ZRC3308-B10 mAb | 4.05E+06 | 6.39E-04 | 1.60E-10 | 2.73E+06 | 3.49E-05 | 1.30E-11 |
| ZRC3308 cocktail (1:1) | 2.93E+06 | 5.41E-04 | 1.85E-10 | 1.47E+06 | 3.73E-05 | 2.60E-11 |
|  | rhFcRn |  |  | rhFcγRIIIa-Phe |  |  |
| Wild type IgG1 | 1.06E+06 | 2.04E-01 | 1.93E-07 | 3.30E+04 | 5.96E-02 | 1.81E-06 |
| ZRC3308-A7 mAb | 1.68E+06 | 2.28E-02 | 1.36E-08 | NB | NB | NB |
| ZRC3308-B10 mAb | 1.01E+06 | 3.42E-02 | 3.40E-08 | NB | NB | NB |

\*NB: No binding

The binding of the mAbs to the RBD of SARS-CoV-2 S1 protein is expected to inhibit the interaction of S1 protein to the ACE-2 receptor. The inhibition of RBD binding to ACE-2 in presence of ZRC-3308 was measured in terms of 50 % inhibitory concentration (IC_50_) by plotting a 4 parameter fit curves of antibody concentration vs. % inhibition as shown in Fig.1i and 1j. Both the mAbs were able to bind to the spike protein RBD and inhibit its binding to ACE2 receptor at sub nM IC_50_ concentrations.

### Binding of ZRC3308 to recombinant human neonatal Fc receptor (rhFcRn), FcγRIIIa-Phe and C1q

ZRC3308 mAbs have been designed to carry mutations in the Fc backbone to improve their FcRn binding and to reduce the binding of antibodies to rhFcγRIIIa-Phe. The ZRC3308-A7 and ZRC3308-B10 antibodies which carry LS mutation in the Fc backbone showed ~10 fold higher affinity for binding to rhFcRn as compared to the wild type IgG1 (Fig.2a-2c, Table 1). The antibodies showed no detectable binding with rhFcγRIIIa-Phe when compared to the wild type IgG1 indicating that ZRC3308-A7 and B10 antibodies are expected to show negligible NK cell mediated effector functions (Fig.2d-2f, Table 1). No binding was observed even at very high mAb concentration of 10μM. C1q binding in terms of effective concentration of mAb leading to 50% maximal binding (EC_50_) was assessed. ZRC3308-A7 (EC_50_=20.55 μg/ml) and ZRC3308-B10 (EC_50_=46.12 μg/ml) showed reduced binding when compared to wild type IgG1 (EC_50_=3.18 μg/ml).

**Fig. 2:**
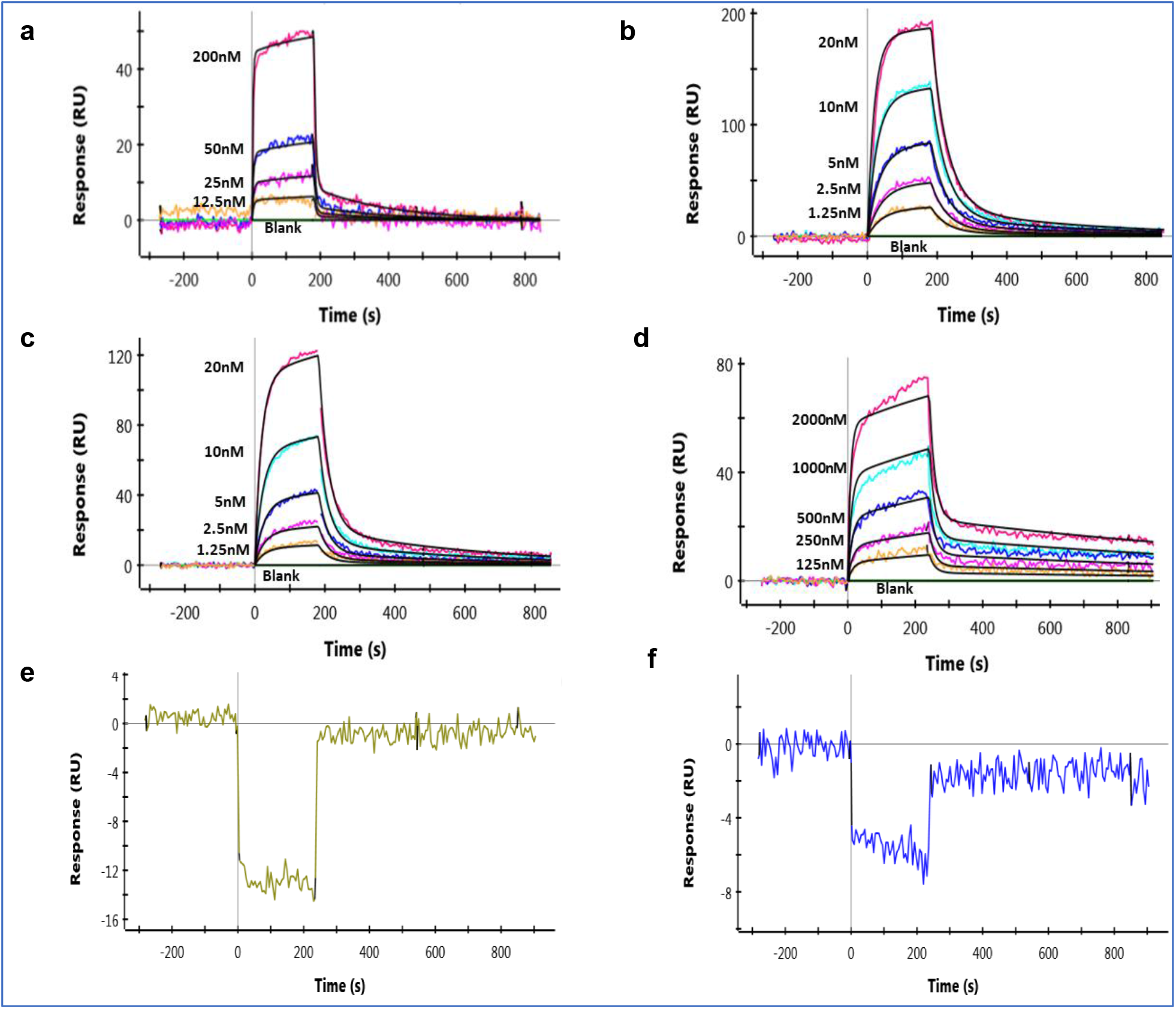
Binding of mAbs to rhFcRn and FcγRIIIa-Phe. Binding sensograms of **a.** wild type IgG1 **b.** ZRC3308-A7 **c.** ZRC3308-B10 to rhFcRn. Binding sensograms of **d.** wild type IgG1 **e.** ZRC3308-A7 and **f.** ZRC3308-B10 to rhFcγRIIIa-Phe.

### Virus neutralizing ability of ZRC3308 mAbs

The neutralizing activities of antibodies were confirmed by both infection of pseudotyped luciferase lentivirus in the HEK293 ACE2 expressing cells and live SARS-CoV-2 in VeroE6/Vero CCL81 cells. The mAbs of the ZRC3308 cocktail exhibited potent neutralization activity against pseudotyped luciferase lentivirus, with an IC_50_ of 8.79 ng/mL and 9.82 ng/mL for ZRC3308-A7 and ZRC-3308-B10, respectively (Fig.3a). The live virus plaque reduction neutralization test (PRNT) using SARS-CoV-2 B.1 variant also showed potent neutralization activity in sub picomolar range with an IC_50_ of 0.1392, 0.1398 and 0.2522 ng/mL for ZRC3308-A7, ZRC-3308-B10 and for the cocktail respectively (Fig.3b). ZRC3308 cocktail showed PRNT_50_ titre of 411781, 327205, 456261, 456261 against B.1.1.7, B.1.351, B.1.617.2 and B.1.617.2 AY.1 variant respectively.

**Fig. 3.**
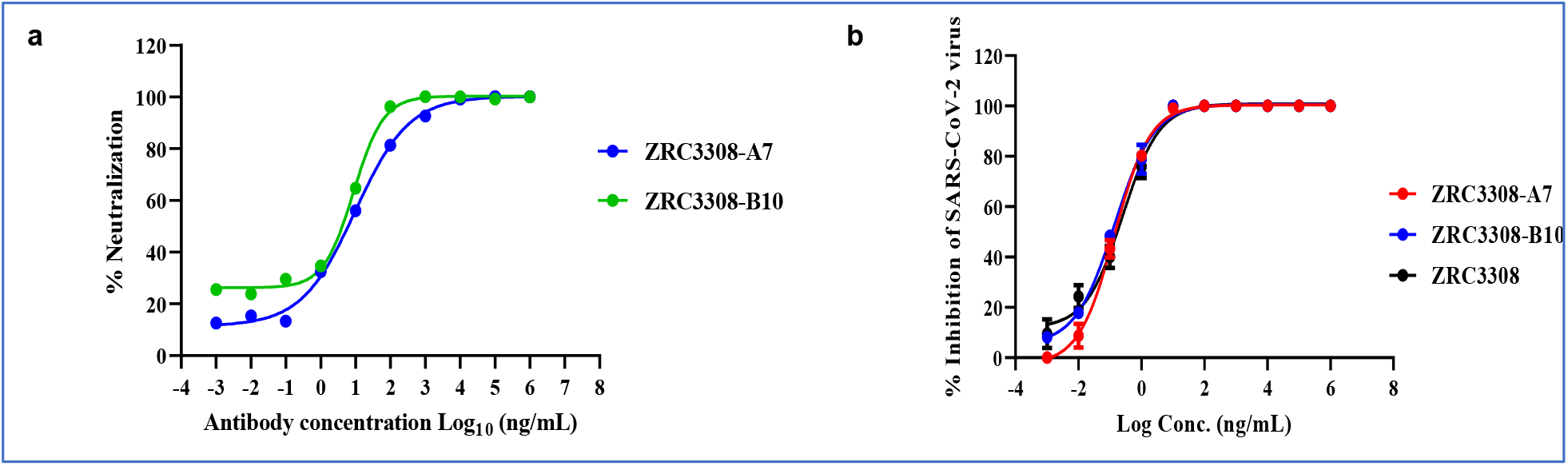
Neutralization potential of mAbs of the ZRC3308 cocktail. **a.** The percent neutralization relative to virus only infection control against the serial dilutions of ZRC3308-A7 and ZRC3308-B10 in the pseudovirus assay. **b.** The percent inhibition of SARS-CoV-2 against serial dilutions of ZRC-3308A7, ZRC-3308B10 and the ZRC3308 cocktail (1:1) antibodies as compared to control without antibodies measured at 72 hours post-infection.

### Pharmacokinetic study in Syrian hamsters

Considering the tissue bioavailability and PRNT50 data, we performed a pharmacokinetic study with doses of 50 mg/kg, 5 mg/kg and 1 mg/kg doses of ZRC3308 cocktail in 1: 1 ratio in Syrian hamsters for a period of 7 days. A dose dependent increase was observed with all the doses of ZRC3308 cocktail in maximum concentration (C_max_), area under curve (AUC) _last_ and AUC _inf-obs_ of the mAb in the serum (Supplementary Table 1). The serum levels of the mAbs remained constant without much reduction till 7 days (Supplementary Figure 1).

### ZRC3308 mAb prophylaxis in Syrian hamsters and virus challenge

For the study, hamsters (n=12/group) were treated with the ZRC3308 cocktail (50 mg/kg, 5mg/kg and 1 mg/kg ZRC3308 cocktail) 48 hours prior to the SARS-CoV-2 infection (Fig. 4a). No clinical signs post was observed following the mAb cocktail treatment and virus infection. Body weight was significantly reduced in the placebo group (mean ± standard deviation (SD) = −7.06 ±3.13 %) compared to 50 mg/kg (mean ± SD = −0.73± 2.1 %, p<0.00005), 5mg/kg (mean ± SD = −1.44 ± 2.42 %, p<0.00005) and 1 mg/kg (mean ± SD = −3.2 ±1.24 %, p <0.0005) treatment groups on 3 days post infection (DPI) (Fig. 4b). On further days even though body weight reduction was there, it was not significant. A sustained level of mAbs in serum without much decrease was observed during the study period (Supplementary Table 2). Viral genomic RNA (gRNA) and subgenomic RNA (sgRNA) levels in the upper and lower respiratory tract showed a decreasing trend in all the mAb treated groups till 7 DPI (Fig.4c-4g). The viral load was significantly reduced in the nasal turbinates (p<0.05) and lungs (p<0.05) in the 50mg/kg dose group on 3, 5 and 7 DPI compared to the placebo group. Other dose groups i.e., 5mg/kg and 1 mg/kg groups also showed a reduction in the viral gRNA load in lungs compared to placebo although not statistically significant. Viral sgRNA could not be detected in the lungs of 50 mg/kg dose group from 3DPI (p<0.05).

**Fig. 4:**
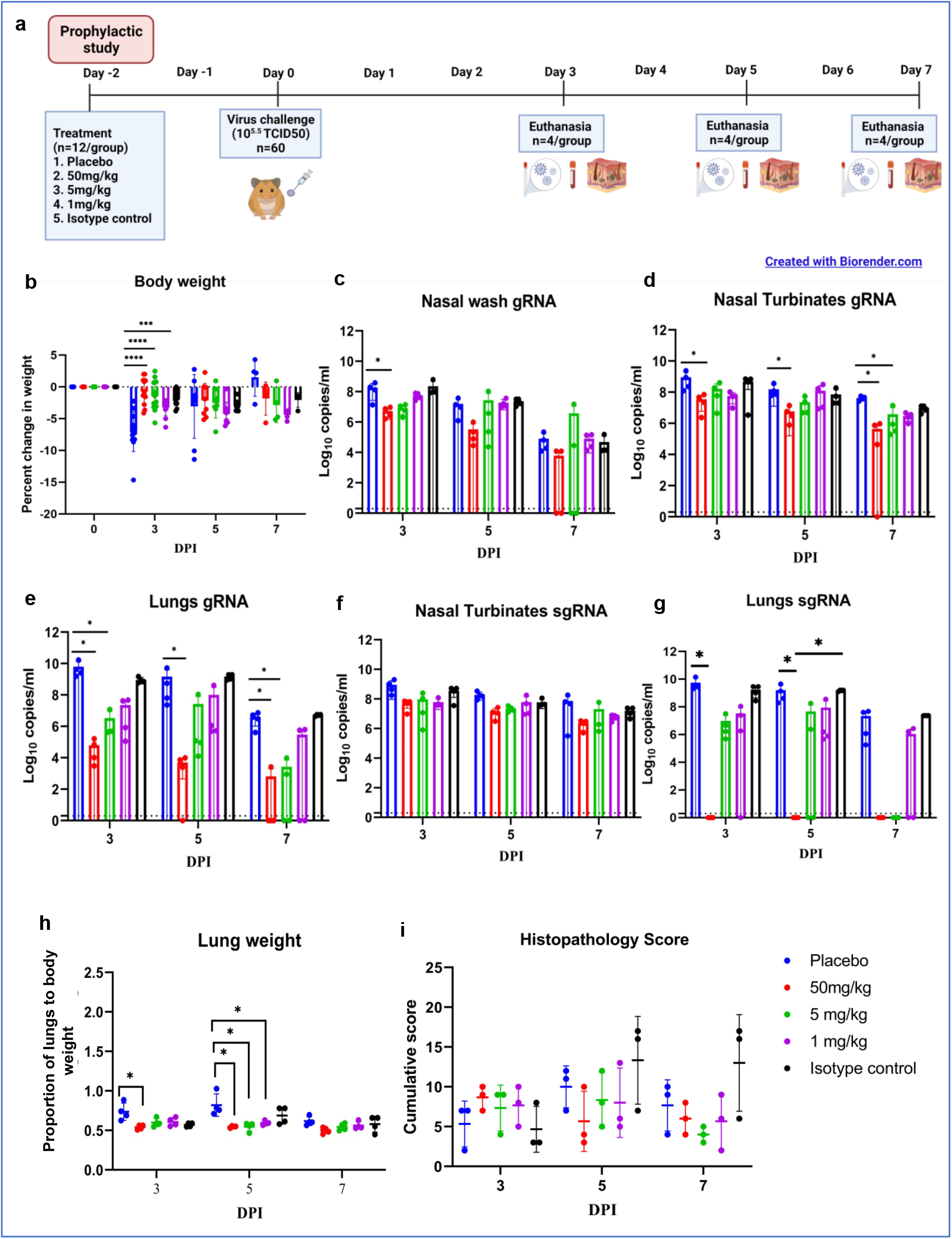
Prophylactic mAb treatment in hamsters. **a**. Study design of the prophylactic study **b.** Percent bodyweight change in hamsters on 3 (n=12/group), 5 (n=8/group) and 7 (n=4/group) DPI. Viral gRNA load in **c.** Nasal wash **d.** Nasal turbinates and **e.** lungs post infection. Viral sgRNA load in **f.** Nasal turbinates and **g.** lungs post infection. **h.** Lungs weight to body weight proportion of hamsters at necropsy. **i.** Cumulative score of lung pathological changes on 3,5 and 7 DPI. Mean ± SD is plotted on the graph and comparison was performed between the treated groups and the control groups. Kruskal–Wallis test followed by Mann–Whitney test was used to assess statistical significance. Asterisk indicates significant difference between the means with ****, ***, * representing p<0.00005, p< 0.0005, p<0.05 respectively and the dotted line on the graph indicates the assay limit of detection.

Grossly, mild congestive changes were observed in lungs in all the mAb treated and the control groups on 3 DPI. On 5th DPI, congestion and hemorrhagic changes were more prominent in the 1 mg/kg dose group, isotype antibody and placebo groups. Lungs of the 50 mg/kg and 5 mg/kg groups appeared normal or with minimal gross changes on 5 and 7 DPI. The proportion of lungs weight to body weight in hamsters was also significantly higher in the placebo group (p<0.05) on 5 DPI compared to the mAb administered group (Fig. 4h). On histopathological examination, the 50 mg/kg and 5 mg/kg dose groups showed mild changes on 3, 5 and 7 DPI (mild alveolar consolidation, focal septal thickening and inflammatory cell infiltration) whereas the 1mg/kg group and placebo group showed moderate lesions (diffuse exudative changes in the alveolar lumen, septal thickening, marked inflammatory cell infiltration in the interstitial space and in the peri bronchial region). The isotype antibody control showed severe pneumonic changes of diffuse involvement compared to other groups (Fig.4e, Fig.5).

**Fig. 5:**
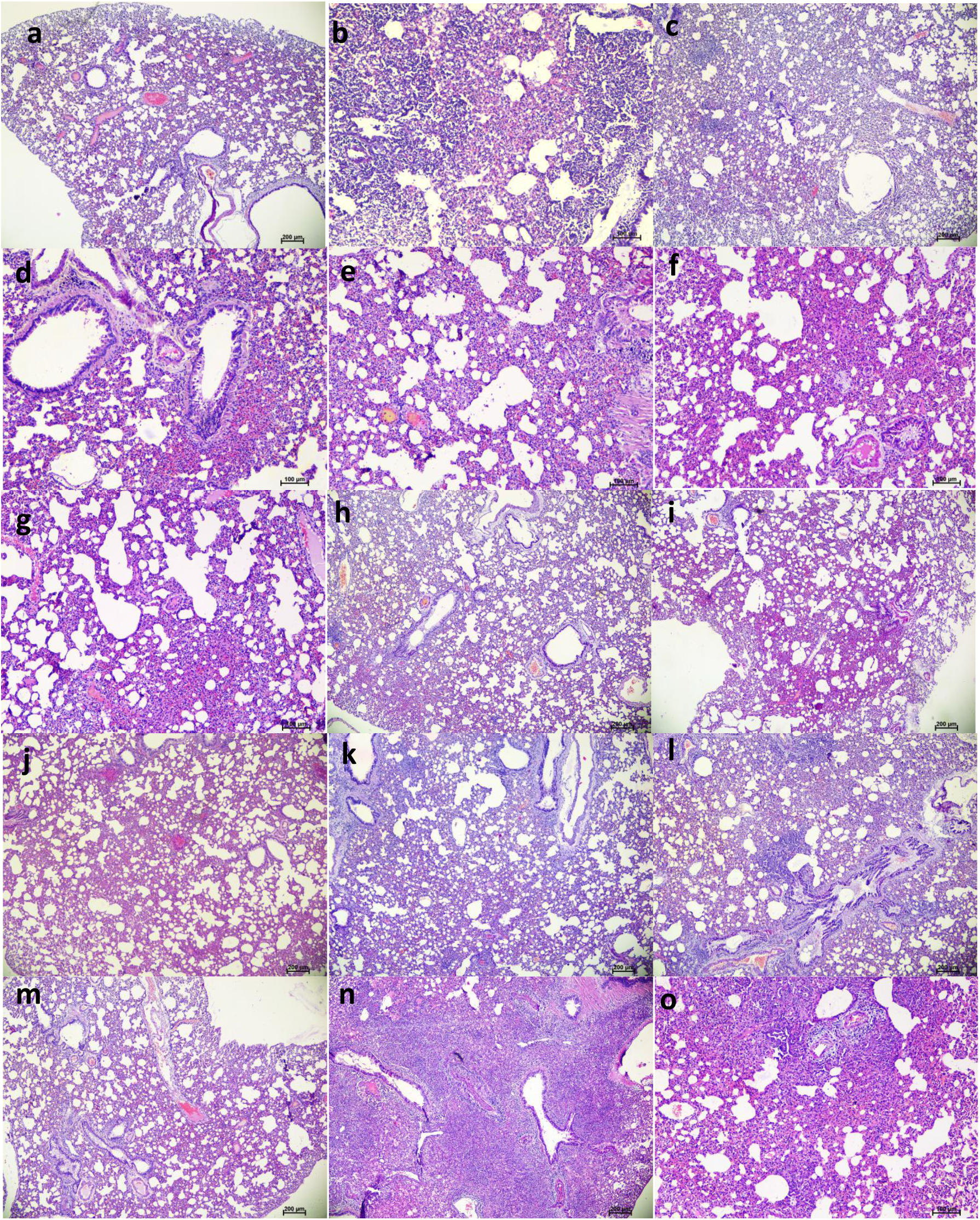
Histopathological observations in lungs of hamsters treated with ZRC3308 cocktail prophylactically. Lungs of placebo group **(a)** on 3DPI showing congestion and focal area of consolidation **(b)** on 5 DPI showing diffuse areas of consolidation, congestion and mononuclear infiltration **(c)** on 7 DPI showing congestion and consolidation. Lungs of 50 mg/kg dose prophylactic group **(d)** on 3DPI showing alveolar capillary congestion **(e)** on 5DPI & **(f)** 7DPI showing alveolar septal thickening and congestive changes. Lungs of 5mg/kg dose group on **(g)** 3 DPI**, (h)** 5 DPI & **(i)** 7 DPI showing congestive changes. Lungs of 1 mg/kg dose group on **(j)** on 3DPI showing congested vessels, **(k)** on 5DPI & **(l)** 7DPI showing congestion and foci of mononuclear cell infiltration in the peri bronchial region. Lungs of the isotype antibody control group on **(m)** 3 DPI showing severe congestion (**n)** on 5DPI showing diffuse pneumonic changes and on 7 DPI showing alveolar septal thickening and consolidation.

### ZRC3308 mAb therapy in Syrian hamsters following SARS-CoV-2 challenge

We studied the therapeutic potential with 10^5.5^ TCID50 (high virus dose) at 24 hours (Fig. 6a) and 6 hours post challenge and with 10 ^3.5^ TCID50 (low) virus dose at 24 hours after infection. In the 10^5.5^ TCID50 challenged group, no significant difference could be observed with body weight loss among groups, whereas gain in body weight was observed in 50mg/kg group (Fig.6b). The mAb levels in the serum remained constant from day 3 to day 7 without much drop (Supplementary Table 2). Although gRNA and sgRNA levels in the respiratory tract showed a decreasing trend in all the groups till 7DPI, there was no statistical significance in comparison to control in the viral load (Fig. 6c-6g). Grossly pneumonic changes were observed in all groups with comparable lung weights to that of placebo group (Fig. 6h). On histopathological examination, moderate to severe pneumonic changes were observed in all treated and control groups (Fig. 6i, Supplementary Figure 1).

**Fig. 6:**
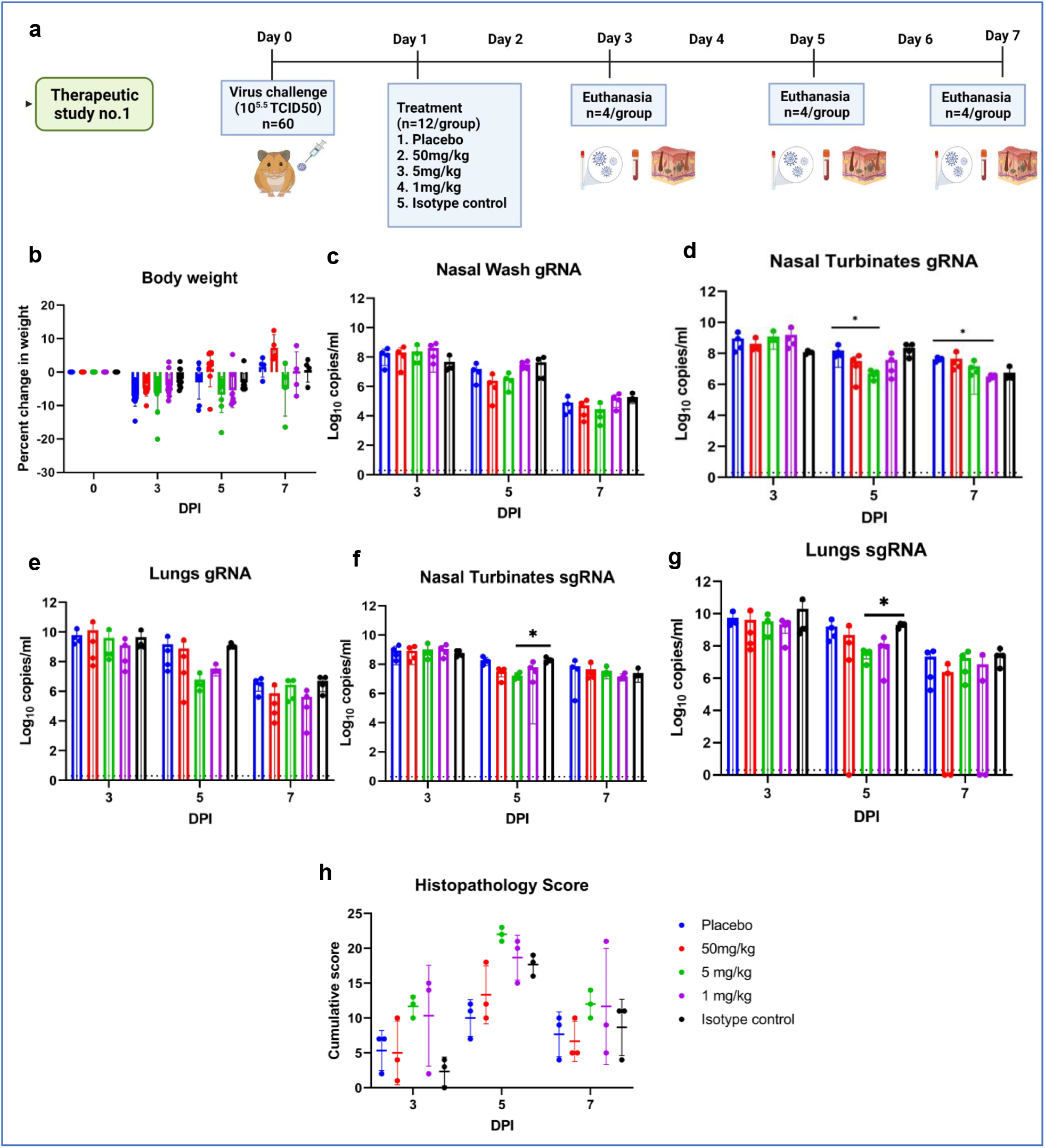
mAb therapy at 24 hours post high virus dose challenge in hamsters. **a.** Study design **b.** Percent bodyweight gain in hamsters on 3 (n=12/group), 5 (n=8/group) and 7 (n=4/group) DPI. Viral genomic RNA load (n=4/group) in (**c)** nasal wash **(d)** nasal turbinates and **(e)** lungs post infection. Viral sub genomic RNA load in hamsters (**f)** nasal turbinates and **(g)** lungs post infection. **h.** Cumulative score of lung pathological changes in hamsters (n= 3/ group) post virus challenge. Mean ± SD is plotted on the graph and comparison was performed between the treated groups and the control groups. Kruskal–Wallis test followed by Mann–Whitney test was used to assess statistical significance. Asterisk indicates significant difference between the means with * representing p<0.05 respectively and the dotted line on the graph indicates the assay limit of detection.

Further we performed therapeutic evaluation of the mAb cocktail (50 mg/kg and 5 mg/kg) at 6 hours post virus (10^5.5^ TCID50) infection to understand the ability of mAb to neutralize at a shorter interval treatment (Fig. 7a). The body weight reduction was observed in both the treated groups (Fig.7b). Although the placebo group showed considerably higher body weight loss than the mAb treated groups, it was not statistically significant. The average gRNA load in nasal wash, nasal turbinate and lungs on 3 and 5 DPI did not show any significant difference among groups (Fig.7c-7e). SgRNA levels showed reduction in comparison to placebo group and were completely cleared from nasal turbinates and lungs by 5DPI in all mAb treated groups (Fig.7f,7g). On histopathological examination, mild changes were seen in the 50 mg/kg dose group whereas the hamsters of 5 mg/kg group showed to mild to moderate changes and the placebo group showed moderate pneumonic changes (Fig 7i, Supplementary Figure 2).

**Fig. 7:**
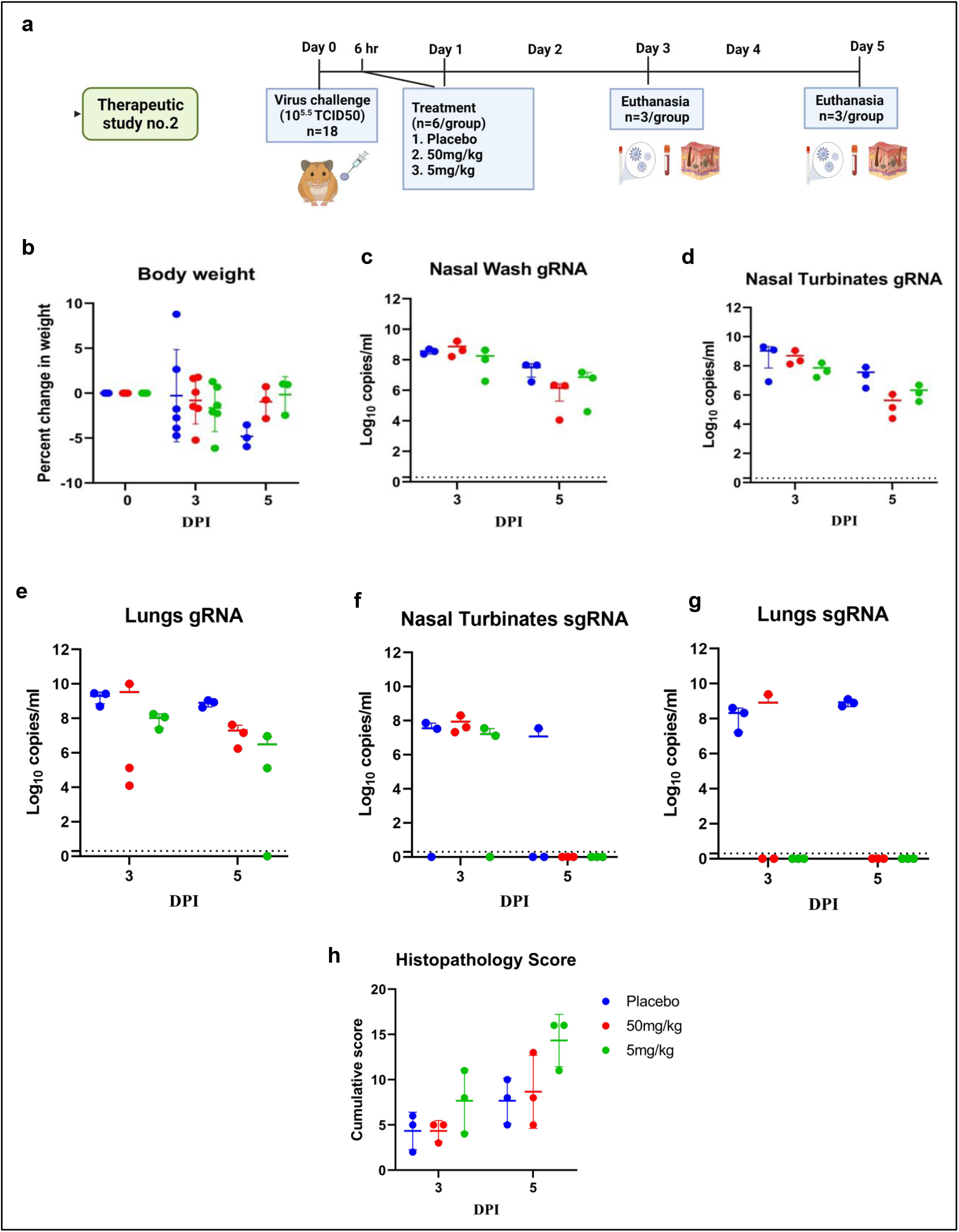
mAb therapy at 6 hours following high virus dose challenge in hamsters. **a.** Study design **b.** Percent bodyweight gain/loss in hamsters on 3 (n= 6/group) and 5 (n=3/group) post SARS-CoV-2 infection. Viral gRNA load (n=3/group) in **(c)** nasal wash **(d)** nasal turbinates and **(e)** lungs in hamsters post virus challenge. Viral sgRNA load (n=3/group) in **(f)** nasal turbinates and **(g)** lungs in hamsters post virus challenge. **h.** Cumulative histopathological score of lung pathological changes in hamsters (n=3/group) on 3 and 5 DPI. Mean ± SD is plotted on the graph and comparison was performed between the treated groups and the control groups. Kruskal-Wallis test followed by Mann-Whitney test was used to assess statistical significance. The dotted line on the graph indicates the assay detection limit.

In the 10^3.5^ TCID50 virus dose group, only minimal bodyweight reduction was seen in the 50 mg/kg and 5 mg/kg mAb treated group (Fig 8a, 8b). The 50 mg/kg group showed consistent viral load reduction on 3 and 5DPI in nasal wash (p <.0.05), lungs (p <.0.05) and nasal turbinates (p <.0.05) (Fig. 8c-e). SgRNA levels in the lungs (p <.0.05) and nasal turbinates (p <.0.05) also were significantly lower compared to placebo group (Fig.8f,8g). On histopathological examination, mild to moderate changes were seen in the 50 mg/kg dose group with a lower histopathological score whereas the hamsters of 5 mg/kg group and placebo group showed comparable disease severity (Fig 8i, Fig 9).

**Fig. 8:**
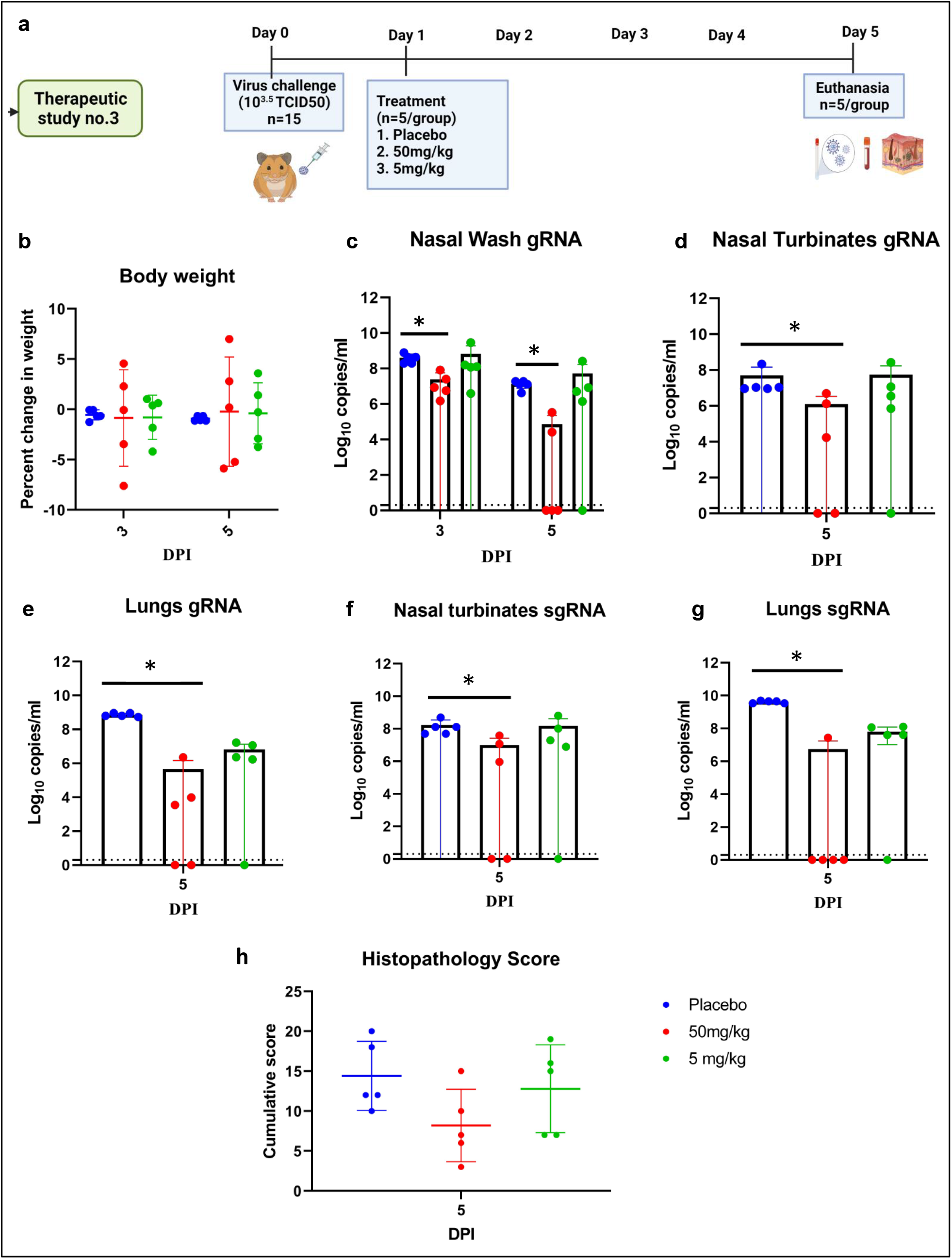
mAb therapy at 24 hours post infection with 10^3.5^ TCID50 virus dose challenge in hamsters. **a.** Therapeutic study design. **b.** Percent bodyweight gain/loss in hamsters on 3 (n= 5/ group) and 5 (n= 5/ group) DPI. Viral gRNA load (n= 5/ group) in **(c)** nasal wash **(d)** Nasal turbinates and **(e)** lungs in hamsters post challenge. Viral sgRNA load (n= 5/ group) in **(f)** Nasal turbinates and **(g)** lungs in hamsters post infection. **h.** Cumulative histopathological score of lung pathological changes in hamsters (n= 5/ group) post virus challenge. Mean ± SD is plotted on the graph and comparison was performed between the treated groups and the control groups. Kruskal–Wallis test followed by Mann–Whitney test was used to assess statistical significance. Asterisk indicates significant difference between the means with * representing p<0.05 and the dotted line on the graph indicates the assay limit of detection.

**Fig. 9.**
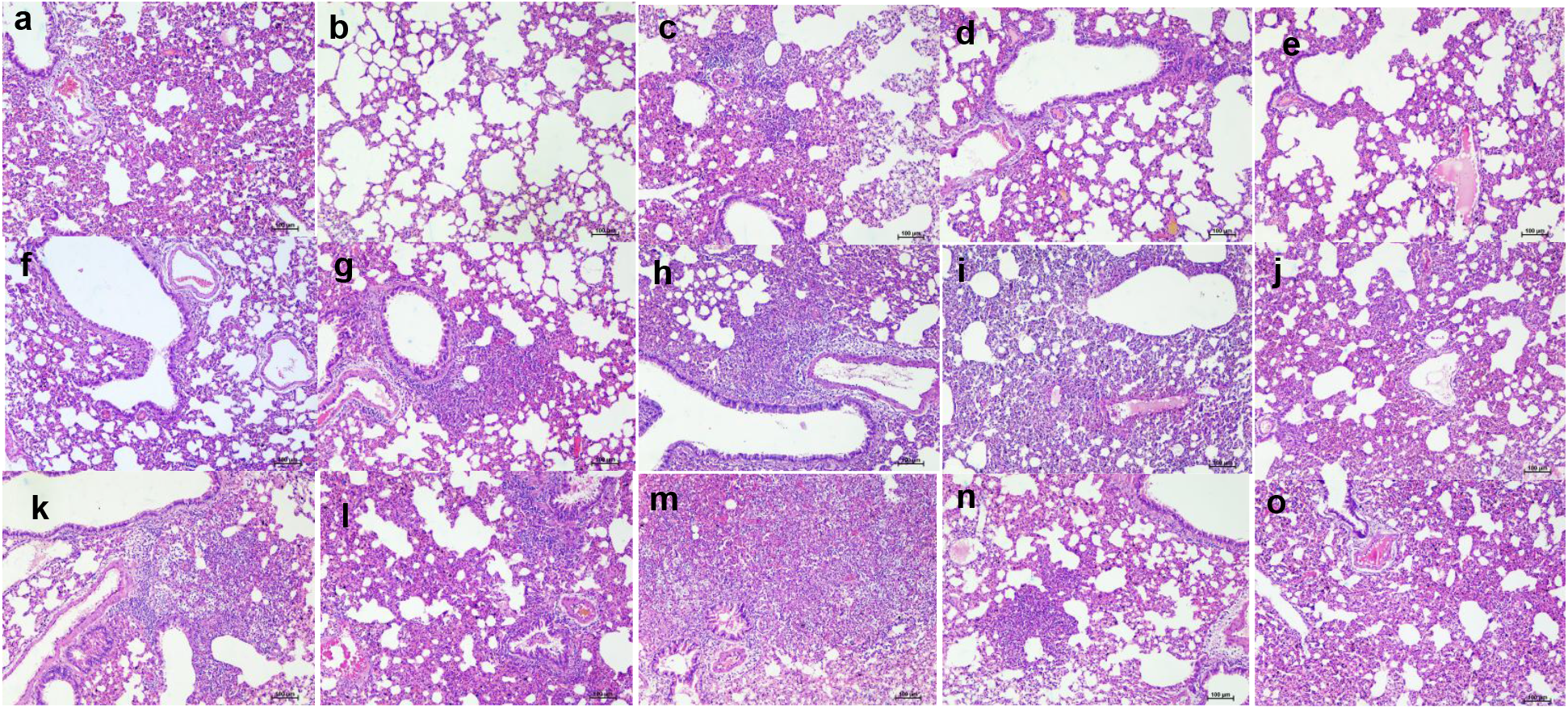
Histopathological changes in lungs of hamsters which received mAb therapy 24 hours post infection with 10^3.5^ TCID50. Lungs of hamsters of 50mg/kg dose group on 5DPI showing **(a)** engorged alveolar capillaries (**b)** emphysematous changes (c) consolidation and mononuclear cell infiltration **(d)** foci of alveolar capillary congestion and **(e)** alveolar septal thickening. Lungs of hamsters of 5mg/kg dose group on 5DPI showing (f) focal areas of congestion and septal thickening (**g)** peribronchial inflammatory cell infiltration and foci of consolidative changes **(h)** diffuse inflammatory cell infiltration in the alveolar paranchyma **(i)** diffuse alveolar changes and congestion and **(j)** diffuse alveolar septal thickening. Lungs of placebo group on 5DPI showing **(k)** diffuse inflammatory cell infiltration in the peribronchial area **(l)** peribronchial inflammatory cell infiltration and alveolar septal thickening **(m)** diffuse alveolar consolidation and inflammatory cell infiltration **(n)** foci of mononuclear cell infiltration and septal thickening **(o)** diffuse alveolar septal thickening **a**nd congestive changes.

## Discussion

COVID-19 pandemic shows no signs of subsiding as evident from the repeated resurgence even after immunization of majority of the population in some countries. Vaccines against SARS-CoV-2 infections has been rolled out as prophylactic interventions but the treatment options are still very limited. mAbs appear to be promising candidates for SARS-CoV-2 as evident by the EUA received by 3 mAb therapies from USFDA which have also been granted approvals in other countries ^13–15^. The ZRC3308 mAbs showed good binding affinity similar to other mAb products for treatment for SARS-CoV-2^14^. As these antibodies bind non-competitively to the RBD of the spike protein and are equipotent in terms of binding to RBD and virus neutralization potential, the 1:1 cocktail diminishes the ability of variants to escape the treatment. *In vitro* PRNT against variants also demonstrated the cross protection. The mutations designed to reduce the immune effector functions of the ZRC3308 mAbs led to a marked reduction in C1q binding indicating reduced complement-dependent cytotoxicity function and no detectable binding to rhFcγRIIIa-Phe further indicating negligible NK cell-mediated antibody-dependent cell cytotoxicity. The increased serum half-life is also reported to be associated with improved lung bioavailability^16^.ZRC3308 mAbs designed to have increased serum half-life effectively brought down sgRNA levels in the lungs of treated animals in both prophylactic and therapeutic settings.

We have used Syrian hamster model to evaluate the protective efficacy of the mAb cocktail. The model has advantages of exhibiting overt clinical signs like body weight loss, replicating to high virus titers in the respiratory tract and severe pneumonia following SARS-CoV-2 infection which can be used as criteria to evaluate countermeasures ^17,18^. Viral load reduction is used as criteria to evaluate the effect of mAb in magnitude of infection in human clinical trials^19,20^. A significant viral load reduction in the upper respiratory tract and lungs of hamsters of the prophylactic group and therapeutic group of hamsters infected with 10^3.5^ TCID50 virus dose. The decrease in viral load was found proportional to the high antibody concentrations. Therapeutic use at 24 hours post infection with a higher virus dose of 10^5.5^ TCID50 could not result in viral gRNA reductions or prevention of pneumonia, whereas treatment at 6 hours could reduce the viral sgRNA loads and body weight loss even though not significant. In an earlier study where the REGN-COV2 prophylactic efficacy was studied in hamsters, no significant reduction could be observed in the lung viral load on 7^th^ day post infection even though there was a decrease in the average viral gRNA load^21^. Although, REGN-COV2 could cause prevention of body weight loss therapeutically at 50mg/kg dose rate in hamsters, the gRNA and sgRNA loads were found non-significant compared to the placebo control. We also observed similar results of no reduction in gRNA load and body weight gain with our mAb cocktail following treatment at 24 hours post infection with high virus dose. When the inoculation dose was lowered, the viral load in the 50mg/kg group was significantly lowered. The reduction in the viral sgRNA in the nasal wash and nasal turbinates in the mAb treated animals is important as upper respiratory tract viral load is a key determinant of transmission.

Wild type IgG1 effectively provides viral clearance but due to their effector functions could increase cytokine release, resulting in pronounced inflammation and tissue destruction. These antibodies by attaching to virus infected cells can mark them for destruction by immune cells. In case of COVID-19, a small delay in treatment may lead to a significantly higher number of host cells getting infected which upon treatment with wild type IgG1 anti-viral mAbs may induce a very strong inflammatory response causing organ damage^8^. It is difficult to demonstrate benefit of mAb in severe COVID-19 cases, where inflammation and coagulopathy induce more damage than viral replication^22^. We also observed the importance of the timing of the mAb treatment in the present study, as similar serum antibody concentration in the blood in the therapeutic group as that of prophylactic group at 3DPI could not prevent the disease progression in the former. However, we have not assessed the lung availability of the mAb cocktail following the post exposure treatment. SARS-CoV-2 produces severe lung disease in hamster model within a short incubation period of 3-5 days^17,18^. The disease severity in hamsters depends on the inoculum dose and the pneumonic changes will set in too rapidly at high virus doses in hamsters making it difficult to demonstrate therapeutic efficacy in the model^21,23,24^. Considering this, we included two more sub studies with shorter time interval for treatment considering the replication cycle length of SARS-CoV-2 and with a lower virus challenge dose. In the both these groups, we observed reduction in viral load following treatment. However, the 6-hour therapeutic group had limitation of small sample size. The interim results of the phase 1-3 clinical trials (ClinicalTrials.gov number: NCT04425629) of the mAb REGN-COV2 in patients with early infection showed a reduced SARS-CoV-2 RNA level in nasopharyngeal swab samples following mAb administration^19^. Similar results were reported in case of bamlanivimab (LY-CoV555) treatment in patients with a median of 4 days after symptom onset^20^. Anti-SARS-CoV-2 mAbs which received EUA like bamlanivimab plus etesevimab and casirivimab plus imdevimab have also not been shown to be beneficial in hospitalized patients with severe COVID-19 and these products warns of worst clinical outcomes in such patients^14,15^. We also observed severe pneumonic changes in the group infected with a high inoculum dose treated with mAb 24 hours post infection. But the similar changes were not observed in the prophylactic and therapeutic groups exposed with a lower virus dose. Rapid induction of pneumonic changes in hamster model is a limitation in the therapeutic evaluation in the present study. The trials have also shown low clinical benefits with LY-CoV555 treatment in hospitalized COVID-19 patients^25^. In the present study, we have characterized ZRC 3308 mAb cocktail for COVID 19 treatment, which was found to be cross neutralizing and a promising candidate for the prophylactic use and for therapy in early cases which have not progressed to severe disease.

## MATERIALS AND METHODS

### Ethics statement

The study was approved by Institute Animal Ethics committee. The study was performed according to the guidelines of Committee for the Purpose of Control and Supervision of Experiments on Animals, Government of India.

### mAb (ZRC3308-A7 and ZRC3308-B10) generation

The codon optimized, light and heavy chain genes of ZRC3308-A7 (CAS RN: 2640223-84-1) and ZRC3308-B10 (CAS RN: 2640224-48-0) were cloned in dual assembly eukaryotic expression vector, DGV-GS, with Glutamine synthase selection marker (DGV-GS). Separate DGV-GS vectors (linearized by PvuI enzyme) encoding the light and heavy chain genes of ZRC3308-A7 and ZRC3308-B10 were used for transfection in CHOK1SVGSKO (Lonza, Switzerland) cell line by electroporation following pre-optimized parameters using Neon transfection system (Invitrogen). Post transfection, the transfectant pools were subjected to selection in the presence of different concentrations (0, 12.5, 25 μM) of Methionine sulphoximine (MSX) (Sigma, USA) in Pro-CHO-5 media (Lonza, Switzerland).

Cells from the high expressing transfection pools of ZRC3308-A7 (25 μM MSX) & ZRC3308-B10 (12.5 μM MSX) were plated over Clona Cell CHO CD medium (StemCellTM Technologies, Canada) supplemented with Clone detect Human IgG H+L specific, fluorescein (Molecular Devices, USA), for clonal selection using ClonePix2 (Molecular Devices, USA). Post, 7-10 days, based on high exterior mean fluorescent intensity, selected clones were aspirated and plated in 96 well plates and expanded further. Pools/clones were subjected to fed-batch cultures to generate material for *in vitro* studies. The cells were seeded (0.3 million/ml) in Acti-Pro production medium (Hyclone, GE, USA) in culti-tubes (TPP, Switzerland) and incubated in a humidified shaker (Kuhner, Switzerland; 37°C, 5 % CO2 and 230 RPM). The cells were cultured for a period of 12-15 days with daily feed of pre-optimized concentration of Cell Boost 7a and 7b supplements (Hyclone, GE, USA) starting from day 3. The antibody concentration in cell culture supernatants were estimated by Protein A high pressure liquid chromatography. Cell culture harvests were purified by Protein A gravity column.

For large scale production of the two mAbs, the above-mentioned process was scaled up to 20 L and then 200 L in suspension bioreactors using a Fed-batch process and utilizing the previously described production media and feeds. The mAbs were purified from the harvested supernatants using a set of purification steps including MabSelect sure (rProtein A) resin (GE life sciences, now Cytivia), a hydrophobic interaction chromatography (GE lifesciences, now Cytivia) and nano-filtration (Merck MilliPore, USA). These purified bulk mAbs were then mixed 1:1 (as per protein content) to generate the cocktail ZRC3308 which was used for all the *in vitro* and *in vivo* studies.

### Surface Plasmon resonance (SPR) analysis

Binding kinetics and affinities towards RBD protein, S Trimer protein, rhFcRn and recombinant human FcγRIIIa-Phe of the individual mAbs ZRC3308-A7, ZRC3308-B10 and cocktail were assessed using Surface Plasmon Resonance technology on a ProteOn XPR 36 instrument (Bio-Rad, USA) using a GLC sensor chip. Filtered and degassed PBST (10mM phosphate buffer, 150mM NaCl, 0.005% Tween 20, pH 7.4) was used as running buffer for all the assays except for rhFcRn binding assay, where PBST buffer with pH 6.0 was used. All the reactions were carried out at 25°C. RBD protein (Cat No: SPD-C82E9; Make: Acro Biosystems, USA) and S Trimer protein (Cat No: SPN-C52H8; Make: Acro Biosystems, USA) were immobilized on the sensor chip surface in two different channels using the standard amine coupling chemistry. To measure the association rate constant (k_assoc_) and dissociation rate constant (k_dissoc_) for the immobilized proteins, five dilutions of mAbs (individual and cocktail) were prepared in PBST running buffer and injected at a flow rate of 50 μL / min with an association time of 300s and dissociation time of 600s. At the end of each cycle, the chip surface was regenerated using 18 s injection of 4M MgCl2. Using the ProteOn Manager Software v 3.1.0.6, the parameters were obtained by fitting the double-reference subtracted data to a 1:1 binding model.

rhFcRn receptor (Cat: CT009-H08H; Sino Biologics, China) was immobilized on a GLC chip using amine coupling chemistry. To measure the k_assoc_ and k_dissoc_, five dilutions of each of the two ZRC3308mAbs were prepared and injected at a flow rate of 100 μL / min with an association time of 180s and dissociation time of 600s. After each sample run, the chip surface was regenerated using PBST, pH 7.4. rhFcγRIIIa-Phe receptor (Cat: 10389-H27H, Sino Biologics, China) was immobilized on a GLC sensor chip surface using a standard amine coupling chemistry. To measure the k_assoc_ and k_dissoc_, five dilutions of each of the two ZRC3308 mAbs were prepared and injected at a flow rate of 100 μL / min with an association time of 240s and dissociation time of 600s. The data in the form of sensograms were analyzed using the data-fitting programs available with the ProteOn system.

### Pair-wise epitope binning

Epitope binning analysis for the two mAbs was carried out using ProteOn XPR36 instrument to confirm that the two antibodies do not compete with each other for binding to RBD and therefore bind to unique epitopes on RBD. The RBD protein (Cat No: SPD-C82E9; Make: Acro Biosystems, USA) was immobilized on a GLC sensor chip surface using standard amine coupling chemistry. 10mM Phosphate Buffered Saline (PBS) was used as the running buffer. Initially ZRC3308-A7 and ZRC3308-B10 were run at saturating concentrations (at 50 nM) in different channels immobilized with RBD. In the next run, ZRC3308-A7 and ZRC3308-B10 were individually run at 50 nM over both ZRC3308-A7and ZRC3308-B10 captured channels and binding patterns were analyzed.

### RBD-ACE2 binding inhibition

The inhibition potential of the mAbs was assessed using a competitive ELISA. Briefly, ACE-2 protein (Acro Biosystems, USA) was coated onto ELISA plates (Greiner, Germany) at 1 μg/mL concentration in PBS and was incubated (for 12-72 hours) at 2-8°C in a humid chamber. The plate was then blocked using 2.0 % solution of BSA (SD Fine Chem, India). Different concentrations of ZRC-3308-A7 (1500 ng/mL to 2.059 ng/mL, 3 fold serial dilutions) and ZRC3308-B10 (800 ng/mL to 3.277 ng/mL, 2.5 fold serial dilutions) were incubated with 100 ng/mL of Biotinylated RBD (Acro Biosystems, USA) on a plate shaker at room temperature for 60 mins to allow the binding of the mAbs to RBD. This mixture was then loaded onto ACE-2 coated plate to allow the unbound RBD to bind to the coated ACE2. Detection was accomplished using peroxidase conjugated streptavidin at 1/50K dilution. TMB was used as a substrate (Sigma Aldrich, USA). Reaction was terminated using 1 N sulfuric acid and the absorbance was read at 450 nm in a multi-mode plate reader (Molecular devices, USA).

### C1q Binding Assay

A chemiluminescence based ELISA method was used to analyze the C1q binding properties of both ZRC3308A7 and ZRC3308-B10. Plates were coated using different optimal concentrations (350 μg/ml to 2.116 μg/ml) of ZRC3308-A7 or ZRC3308-B10 or wildtype IgG1 (60 μg/ml to 0.363 μg/ml) and was incubated in 37 + 2 °C in incubator for 2 hours. The coated plates were then blocked using 2% skimmed milk (Hi Media, India). This was followed by addition of 8 μg/ml C1q (Sigma, USA) to the plate wells. Detection was accomplished using 1/1000 diluted peroxidase conjugated sheep anti-C1q polyclonal antibody (Abcam, UK). Subsequently Femto substrate (Thermo Scientific, USA) was added, and post 5 minutes incubation in dark, luminescent signals were quantified using multimode reader (Molecular devices, USA) in Luminescence mode. The 4-parameter fit was used to determine the EC_50_ concentrations.

### Pseudovirus based neutralization Assay

HEK 293 ACE2 expressing cells (Scripps institute) were used for the assay. Pseudovirus (an HIV-based luciferase expressing lentivirus pseudotyped with SARS-CoV-2 full length S protein) was obtained from Creative Biogene. One step luciferase assay kit from BPS Bioscience was used for detection. The IC_50_ was defined as the dilution of serum at which the relative light units (RLUs) were reduced by 50% compared with the virus control after subtraction of the background RLUs of the cell control.

In brief, the antibody preparations were diluted to a starting concentration of 1 mg/mL, and serially diluted with 10-fold dilutions to generate a concentration range of 1×10^6^ ng/mL to 0.001 ng/mL. The pseudovirions were diluted in assay medium to achieve a MOI of 50 per well. Pseudovirions were added to the serially diluted antibody preparations and the plate was incubated for 60 minutes at 37°C in a 5% CO2 incubator. Simultaneously, HEK 293 ACE2 expressing cells were trypsinized and counted. HEK 293 ACE2 expressing cells (1.0×10^4^ per well) were seeded in a flat tissue culture plate. After completion of incubation, the mixture containing antibody and pseudovirions, was added to the seeded HEK 293 ACE2 expressing cells and the plate incubated for 24 hrs at 37°C in a 5% CO2 incubator. Following 24 h of incubation the virus containing media was replaced with fresh growth medium and the plate was further incubated for 24-48 hrs at 37°C in a 5% CO2 incubator. The plate was observed for confluency and after reaching 85-90 % confluency it was removed to measure the luminescence. 100 μL of firefly luciferase substrate was added to all the wells and the plate was incubated on shaking condition at room temperature for 30 minutes to check the transduction efficacy. The plate was read using a luminescence microplate reader (SpectraMax^®^ i3x). The obtained data is normalized against the cell control well. The percent neutralization data in relation to the virus only infection control is plotted in a graph against the concentrations of mAbs to obtain the IC_50_ values for the samples. The IC_50_ values were calculated using GraphPad Prism. (GraphPad Prism 8, Inc., San Diego, CA, USA).

### SARS-CoV-2 live virus plaque reduction neutralization test

SARS-CoV-2 isolate belonging to B.1 lineage was used. Vero E6 cells (1.0×10^6^ per well) were seeded to 24-well plates in maintenance medium for 24 hrs at 37°C in a 5% CO2 incubator. Next day mAbs were serially diluted in assay medium (range of 1×10^6^ ng/mL to 0.001 ng/mL). SARS-CoV-2 virus was added to each dilution at 0.01 MOI except the cell controls. The mixtures of virus and antibody were incubated for 60 min at 37°C in a 5% CO2 incubator. Following completion of incubation, the virus and antibody mixtures were added to the pre-seeded Vero E6 cells by first discarding the supernatant followed by replacement with medium containing 2% carboxy methylcellulose. The plates were incubated for a further 72 hours in a CO_2_ humidified incubator at 37°C. Post incubation the cells were fixed with 4% formaldehyde and the plaques were enumerated by staining with crystal violet stain. The number of plaques were counted and the percentage inhibition was calculated in comparison to the number of plaques obtained in the wells that contained only the virus but no antibody (positive control). IC_50_ was calculated using GraphPad Prism 8 (GraphPad Software, Inc., San Diego, CA, USA). Further we performed PRNT_50_ using SARS-CoV-2 variants like B.1.1.7, B.1.351, B.1.617.2 and B.1.617.2 AY.1 as described earlier^26^.

### Pharmacokinetic study in hamsters

A total of 18 female hamsters, aged 11-12 weeks were divided into four groups according to the dose of the cocktail i.e., 50 mg/kg (25 mg/kg of ZRC3308-A7 + 25 mg/kg of ZRC-3308-B10), 5 mg/kg (2.5 mg/kg of ZRC3308-A7 + 2.5 mg/kg of ZRC3308-B10), 1.0 mg/kg (0.5 mg/kg of ZRC3308-A7 + 0.5 mg/kg of ZRC3308-B10), and placebo consisting of 5 animals each except for the placebo which consisted of only 3 animals. Cocktails of ZRC3308-A7 and ZRC3308-B10 were administered via intraperitoneal (I.P.) route and the total duration of the study was 7 days. Blood was withdrawn from the animals before administration of mAb and at 24, 72, 120 and 168 hrs post administration. The serum was separated to assess the concentration of each of the two mAbs by ELISA. For the placebo group, samples were collected only before administration and at 168 hrs.

### ELISA based detection of mAb levels in serum

Two separate ELISA methods were used to detect ZRC3308-A7 and ZRC3308-B10 antibodies. In both the immunoassays, coating reagent used was SARS-CoV-2 S1 protein (Acro Biosystems, USA). Post 24-72 hours of coating and incubation in humidified chamber at 2-8°C, ELISA plates (Greiner, Germany) were blocked using 2% BSA (SD Fine Chem). Subsequently, the calibration curve (ranging from 25 ng/mL to 0.195 ng/mL) was prepared separately for ZRC3308-A7 and ZRC3308-B10 with pooled, naive, hamster serum diluted 500 times in 0.1% BSA in PBST and the serum samples containing ZRC-3308 cocktail were added at appropriate dilutions. Detection was accomplished using 1/100,000 diluted peroxidase conjugated goat anti-human lambda light chain secondary antibody for ZRC3308-A7 (Novus Biologics, USA) and 1/135,000 diluted Goat anti-Human kappa light chain secondary antibody for ZRC3308-B10 (Novus Biologics, USA) to specifically detect the two antibodies. TMB was used as a substrate. The reaction was stopped using 1N sulfuric acid and the ELISA plates were read in multi-mode reader (Molecular devices, USA) at 450 nm.

### Challenge study in Syrian hamsters

A total of 60 female, Syrian hamsters, aged 7-10 weeks were used for the prophylactic study, which were divided into 5 groups of 12 animals each. The groups were high (50mg/kg), medium (5 mg/kg), low (1mg/kg) dose of mAb cocktail, an IgG1 isotype antibody (Trastuzumab @ 50 mg/kg) control and a placebo control. The animals were intranasally inoculated with 0.1 ml of 10^5.5^ TCID50 of SARS-CoV-2 strain isolate belonging to B.1 lineage. The cocktail of mAbs were administered by intraperitoneal injection 2 days before virus infection for the prophylactic group. To the placebo group, only SARS-CoV-2 intranasal virus challenge was performed without any antibody treatment.

There were three sub-studies in the therapeutic arm. For the first therapeutic study, 60 female hamsters aged 7-10 weeks were used which were divided into 5 groups as mentioned for prophylactic study. The mAb treatment was given at 24 hours post infection. The animals were closely observed for any mortality or clinical signs post administration like activity, ruffled fur, discharges from natural orifices, labored breathing and body weight changes. Four animals each of the prophylactic and therapeutic study group were sacrificed by overdose of isoflurane anesthesia on 3, 5 and 7 DPI and necropsy was performed to collect nasal wash, nasal turbinates, lungs and blood samples. Serum was separated and stored at −20°C to test for the level of mAb levels. The organ samples were stored at −80°C for viral load estimation and in 10% neutral buffered formalin for histopathological analysis. The proportion of the lungs to body weight ratio was also determined.

In the second study of therapeutic arm, 18 female hamsters aged 8-10 weeks were used and mAb treatment was given 6 hours post infection with a virus challenge dose of 10^5.5^ TCID50. The 50 mg/kg and 5 mg/kg mAb dose groups along with placebo group with 6 animals each were included in the study. On day 3 and 5, 3 hamsters form each group were sacrificed to assess collect organ and nasal wash samples.

In the third therapeutic study, a challenge virus dose of 10^3.5^ TCID50 was given followed by mAb treatment at 24 hours post infection. Fifteen female hamsters aged 16-18 weeks were used for the study. The 50 mg/kg and 5 mg/kg mAb dose groups along with a placebo group with 5 animals each were included in the study. The hamsters were sacrificed on day 5 to assess lung pathology and viral load in organs.

### Viral RNA quantification

Nasal wash samples collected in 1ml viral transport medium and weighed organ samples homogenized in 1 ml media, were used for RNA extraction. MagMAX™ Viral/Pathogen Nucleic Acid Isolation Kit was used as per the manufacturer’s instructions. E gene Real-time RT-PCR was performed using published primers^27^. SgRNA levels were also estimated by primers targeting N gene of SARS-CoV-2 using published primers^28^.

### Histopathological examination

Lungs samples collected were fixed in 10% neutral buffered formalin and were further processed using routine histopathological techniques. The lesions were blindly scored based on the vascular changes, bronchial lesions, alveolar pathological changes like septal thickening, pneumocyte hyperplasia, consolidation, edematous changes and inflammatory cell infiltration. Each lesion was scored on a scale from 0 to 4 and the cumulative score were represented in a graph.

### Statistical analysis

Graphpad Prism version 8.4.3 software was used for data analysis of the experimental challenge study. Kruskal-Wallis test followed by Mann-Whitney test was used. P < 0.05 were considered to be statistically significant.

## Notes

### Competing Interest Statement

The authors have declared no competing interest.

