## Supplemental data for "ZRC3308 monoclonal antibody cocktail shows protective efficacy in Syrian hamsters against SARS-CoV-2 infection"

**Supplementary data**

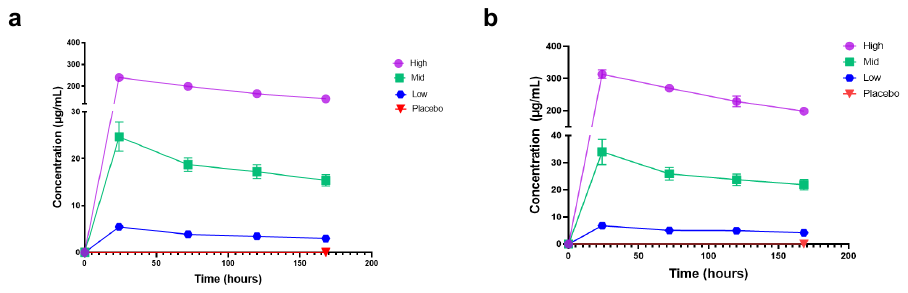

**Supplementary Figure 1: Pharmacokinetic study of ZRC3308 cocktail in Syrian hamsters. a.** Pharmacokinetic profile of ZRC3308-A7 in hamster serum for the 50 mg/kg, 5 mg/kg, 1 mg/kg and placebo. **b.** Pharmacokinetic profile of ZRC3308-B10 in hamster serum for the 50 mg/kg, 5 mg/kg, 1 mg/kg and placebo.

**Supplementary Figure 2: Histopathological changes in lungs of hamsters which received mAb therapy 24 hours post infection.** Lungs of placebo group **(a)** on 3DPI showing congestion and focal area of consolidation, **(b)** on 5 DPI showing extensive areas of congestion septal thickening and **(c)** on 7 DPI showing consolidation and congestion. Lungs of 50 mg/kg dose prophylactic group **(d)** on 3DPI showing congestion and focal area of infiltration **(e)** on 5DPI showing diffuse mononuclear infiltration, pneumocyte hyperplasia and exudative changes and **(f)** on 7DPI showing peri bronchial mononuclear infiltration and haemorrhages. Lungs of 5mg/kg dose group on **(g)** 3 DPI showing haemorrhages, **(h)** 5 DPI showing alveolar exudative changes, multifocal areas of mononuclear infiltration and **(i)** on 7 DPI showing diffuse mononuclear infiltration. Lungs of 1 mg/kg dose group on **(j)** on 3DPI showing severely congested vessels, **(k)** on 5DPI showing congestion and foci alveolar septal thickening with pneumocyte hyperplasia. and **(l)** 7DPI showing diffuse alveolar damage. Lungs of the isotype antibody control group on **(m)** 3 DPI showing severe congestion **(n)** on 5DPI showing peri bronchial mononuclear infiltration and on **(o)** 7 DPI showing infiltration in the peri bronchial area and collapse of surrounding alveoli.

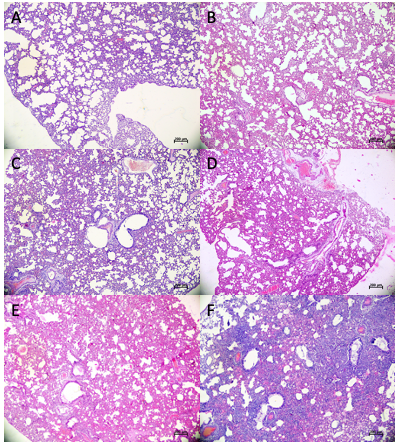

**Supplementary figure 3: Histopathological changes in lungs of hamsters which received mAb therapy 6 hours post infection.** Lungs of 50mg/kg dose group **(a)** on 3DPI showing mild congestion **(b)** on 5 DPI showing congestion and atelectasis. Lungs of 5 mg/kg dose group **(c)** on 3DPI showing congestion and focal area of infiltration and **(d)** on 5DPI showing severe congestion, consolidative changes and focal mononuclear infiltration. Lungs of placebo group showing **(e)** haemorrhage, congestion and exudative changes on 3DPI and **(f)** on 5 DPI showing diffuse mononuclear infiltration, congestion, pneumocyte hyperplasia and consolidation.

**Supplementary Table 1. Pharmacokinetic parameters of ZRC3308 cocktail.** Thepharmacokinetic profile of ZRC-3308A7 and ZRC-3308B10 mAbs in the hamster serum at T_max_ of 24 hours.

| **Dose (mg/kg)** |  | **ZRC3308-A7** | | | **ZRC3308-B10** | | |
| --- | --- | --- | --- | --- | --- | --- | --- |
|  |  | **C _max_ (µg/mL)** | **AUC _last_**  **(h*µ/mL)** | **AUC _ifn_obs_**  **(h*µg/mL)** | **C _max_ (µg/mL)** | **AUC _last_**  **(h*µ/mL)** | **AUC _ifn_obs_**  **(h*µg/mL)** |
| **0.5** | N | 4 | 4 | 4 | 4 | 4 | 4 |
|  | Mean | 5.47 | 619.29 | 1470.99 | 6.82 | 828.74 | 2226.37 |
|  | SD | 0.54 | 29.84 | 187.91 | 0.76 | 42.72 | 286.69 |
| **2.5** | N | 5 | 5 | 5 | 5 | 5 | 5 |
|  | Mean | 24.67 | 2983.02 | 8521.71 | 33.99 | 4130.48 | 14129.56 |
|  | SD | 3.11 | 234.55 | 1714.57 | 4.67 | 360.43 | 3277.83 |
| **25** | N | 5 | 5 | 5 | 5 | 5 | 5 |
|  | Mean | 240.4 | 29624.06 | 72933.74 | 313.48 | 39974.5 | 103423.03 |
|  | SD | 7.19 | 1258.91 | 13487.96 | 13.34 | 1792.82 | 13754.51 |

*T _max_ reported as median

^#^ One animal in 0.5 mg/kg has been excluded from analysis as no drug could be detected which may be due to dosing error

**Supplementary table 2: Serum concentration of monoclonal antibody post virus infection at 3, 5 and 7 days**

| **Group** | **Days post infection** | **50mg/kg dose** | | **5 mg/kg dose** | | **1 mg/kg dose** | |
| --- | --- | --- | --- | --- | --- | --- | --- |
|  |  | **ZRC-3308- A7 Concentration (mg/ml)** | **ZRC-3308- B10 Concentration (mg/ml)** | **ZRC-3308- A7 Concentration (mg/ml)** | **ZRC-3308- B10 Concentration (mg/ml)** | **ZRC-3308- A7 Concentration (mg/ml)** | **ZRC-3308- B10 Concentration (mg/ml)** |
| **Prophylactic** | **3** | 0.461 | 0.616 | 0.133 | 0.117 | 0.005 | 0.002 |
|  |  | 0.368 | 0.422 | 0.139 | 0.127 | 0.006 | 0.002 |
|  |  | 0.372 | 0.435 | 0.14 | 0.129 | 0.006 | 0.002 |
|  |  | 0.414 | 0.515 | 0.136 | 0.123 | 0.005 | 0.002 |
|  | **5** | 0.344 | 0.386 | 0.124 | 0.101 | 0.004 | 0.002 |
|  |  | 0.343 | 0.386 | 0.122 | 0.098 | 0.004 | 0.002 |
|  |  | 0.405 | 0.507 | 0.111 | 0.076 | 0.004 | 0.002 |
|  |  | 0.369 | 0.443 | 0.122 | 0.1 | 0.004 | 0.002 |
|  | **7** | 0.352 | 0.424 | 0.122 | 0.104 | 0.004 | 0.002 |
|  |  | 0.369 | 0.448 | 0.12 | 0.108 | 0.004 | 0.002 |
|  |  | 0.351 | 0.415 | 0.117 | 0.087 | 0.004 | 0.002 |
|  |  | 0.397 | 0.512 | 0.119 | 0.101 | 0.003 | 0.002 |
| **Therapeutic (24 hours)** | **3** | 0.538 | 0.81 | 0.182 | 0.206 | 0.009 | 0.015 |
|  |  | 0.366 | 0.418 | 0.146 | 0.139 | 0.008 | 0.012 |
|  |  | 0.469 | 0.674 | 0.151 | 0.149 | 0.003 | 0.002 |
|  |  | 0.494 | 0.669 | 0.141 | 0.132 | 0.008 | 0.011 |
|  | **5** | 0.472 | 0.625 | 0.141 | 0.132 | 0.006 | 0.008 |
|  |  | 0.512 | 0.722 | 0.13 | 0.11 | 0.006 | 0.009 |
|  |  | 0.507 | 0.718 | 0.137 | 0.126 | 0.005 | 0.008 |
|  |  | 0.223 | 0.155 | 0.141 | 0.134 | 0.006 | 0.009 |
|  | **7** | 0.391 | 0.476 | 0.114 | 0.092 | 0.004 | 0.003 |
|  |  | 0.223 | 0.161 | 0.119 | 0.1 | 0.004 | 0.003 |
|  |  | 0.456 | 0.603 | 0.128 | 0.123 | 0.004 | 0.003 |
|  |  | 0.431 | 0.614 | 0.118 | 0.098 | 0.004 | 0.004 |
| **Therapeutic (6 hours)** | **3** | 0.461 | 0.616 | 0.133 | 0.117 |  |  |
|  |  | 0.368 | 0.422 | 0.139 | 0.127 |  |  |
|  |  | 0.372 | 0.435 | 0.14 | 0.129 |  |  |
|  | **5** | 0.344 | 0.386 | 0.124 | 0.101 |  |  |
|  |  | 0.343 | 0.386 | 0.122 | 0.098 |  |  |
|  |  | 0.405 | 0.507 | 0.111 | 0.076 |  |  |
